# Spatial Uncoupling of Protein Synthesis from Viral Factories during Rabies Virus Infection

**DOI:** 10.64898/2026.10.05.756732

**Authors:** Léonardon Benjamin, Pinto De Magalhaes Boris, Nevers Quentin, Gaudin Yves, Cécile Lagaudrière-gesbert

**Affiliations:** Institute for Integrative Biology of the Cell (I2BC), CEA, CNRS, Université Paris-Saclay, Gif-sur-Yvette, France; Unit Molecular Immunology and Virology (VIM), INRAE, Jouy-en-Josas, France

## Abstract

Replication and transcription of *Mononegavirales* (MNV) occur in cytoplasmic viral factories. In rabies virus (RABV) infected cells, these membrane-less, liquid-liquid phase-separated organelles are called Negri bodies (NBs). Their accumulation of viral mRNAs suggests that, in addition to supporting viral RNA synthesis, they may also serve as sites of viral protein translation. Using immunofluorescence and confocal microscopy, we demonstrate that ribosomal subunit proteins and translation initiation factors are excluded from NBs, including the ribosomal protein RPL40, a component of the large ribosomal subunit involved in the unconventional translation initiation mechanism of RABV. To investigate the spatiotemporal dynamics of protein synthesis during RABV infection, we performed pulse-labeling experiments with the methionine analogue homopropargylglycine (HPG). HPG labeling revealed a rapid accumulation of newly synthesized proteins within NBs, occurring independently of both microtubule and actin cytoskeletons. RABV infection also triggered the formation of stress granules (SGs), in which some ribosomal proteins and translation initiation factors are concentrated. Notably, a marked inhibition of protein synthesis was observed in SG-positive cells. Together, these findings indicate that in RABV infected cells translation occurs outside NBs, followed by the targeting of nascent viral proteins to these compartments.

## Introduction

Viruses are obligate intracellular parasites that entirely rely on the host translational machinery for their replication. They employ specialized strategies to either prevail over cellular transcripts or hijack translation factors and ribosomes for their own mRNA, by frequently using unconventional translation mechanisms [1]. Viruses not only need access to the host translation machinery but must also counteract host innate defenses designed to impair the protein synthesis [2]. Stress granules (SG) formation is one of the key mechanisms to prevent translation of potentially harmful mRNA, such as viral RNA [3]. They are membrane-less organelles that accumulate stalled initiation complexes containing messenger ribonucleoproteins (mRNPs), mRNA, 40S subunits, initiation factors, and RNA-binding proteins (RBPs) [4]. Viral infections can either induce or suppress SG assembly. When induced, SGs often appear only transiently before being disassembled, altered in composition, or hijacked by the virus to support its own replication [5].

Rabies virus (*Mononegavirales* order, *Rhabdoviridae* family, *Lyssavirus* genus) is the causing agent of rabies, a progressive and invariably fatal encephalitis without timely post-exposure treatment [6]. It is an enveloped virus that contains a ∼12 kb non-segmented, negative-sense single-stranded RNA genome. The genome is tightly encapsidated by nucleoprotein N associated with the RNA-dependent RNA polymerase (L) and its cofactor, the phosphoprotein (P), to form the ribonucleoprotein (RNP), which is surrounded by a lipid envelope displaying surface glycoprotein G trimers that mediate host cell attachment and membrane fusion [7]. Following RABV entry, the RNP is released into the cytoplasm and serves as a template for transcription that is catalyzed by the L-P polymerase complex. During transcription, a positive-stranded leader RNA and five capped and polyadenylated mRNAs are synthesized. The replication process yields nucleocapsids containing full-length antigenomic RNA, which in turn serve as templates for the synthesis of genomic RNA. Replication strictly depends upon ongoing protein synthesis to provide the N protein necessary to encapsidate nascent antigenomes and genomes.

A hallmark of RABV infection is the formation of cytoplasmic inclusion bodies known as Negri bodies (NBs), first described by Adelchi Negri in 1903, and long been used as diagnostic markers visible in histopathological specimens from infected brain tissue. Functional characterization of NBs showed that they concentrate viral genomic and antigenomic RNAs as well as all viral mRNAs, and that they were the sites of the synthesis of these viral RNA species [8]. Later, it was shown that these viral factories exhibit properties of liquid organelles formed through liquid-liquid phase separation (LLPS) [9], which was subsequently demonstrated for the viral factories of several other *Mononegavirales* (MNV) [10–14].

With regard to viral protein synthesis, an unconventional translation initiation mechanism requiring the large ribosomal protein RPL40 was characterized for vesicular stomatitis virus mRNAs and was also found to be conserved in other mononegaviruses, including RABV [15]. However, the precise location at which viral mRNA translation occurs is still undefined. Whether RABV translation occurs in NBs alongside the viral transcription machinery or employs another strategy, with replication concentrated in NBs and translation occurring in the surrounding cytoplasm or at NBs periphery, remains an open question.

In this study, we employed ribopuromycylation (RPM) and pulse-labeling with the methionine analog homopropargylglycine (HPG) to map the spatiotemporal dynamics of protein synthesis during RABV infection. By combining these approaches with immunofluorescence analysis of translation machinery components, we demonstrated that viral translation occurs outside viral factories, while *de novo*-synthesized proteins rapidly accumulate within NBs via a cytoskeleton-independent mechanism. Additionally, our findings revealed the recruitment of components of the translational preinitiation complex into stress granules in RABV-infected cells.

## Materials and Methods

### Cell lines and virus

HEK293T cells (human embryonic kidney, ATCC #CRL-3216) and U-373MG cells (human glioblastoma astrocytoma) were purchased from ATCC organization (http://www.lgcstandards-atcc.org). BSR cells, a clone from BHK 21 (Baby Hamster Kidney) were obtained from Dr A. Flamand (Former Laboratoire de Génétique des Virus, Gif-sur-Yvette, France). BSR-T7/5 cell line is a clone of BHK 21 constitutively expressing the T7 RNA polymerase. All these cells were grown in Dulbecco’s modified eagle medium (DMEM) (Gibco-Life Technologies) supplemented with 10% fetal calf serum (FCS) (Gibco-Life Technologies) and 100 units/mL of Penicillin and 100 mg/mL of Streptomycin (AB 1X) (Gibco-Life Technologies). BSR-T7/5 cells were grown in the same medium supplemented with 1 mg/mL Geneticin (Gibco-Life Technologies). The SAD L16-B19 strain of rabies virus (called SAD all along the text) was grown on BSR cells.

## Construction of Recombinant Virus SAD-P-mCherry

The full-length genomic plasmid pSAD L16-B19 was digested with NcoI and PspXI restriction enzymes (New England Biolabs). Two overlapping fragments were amplified by PCR: the first one going from the NcoI site within the P gene to genomic position 3096 and included the mCherry coding sequence fused in-frame to the C terminus of the P coding sequence, and the second fragment going from genomic position 3079 to the PspXI site in G coding sequence. The PCR products and the digested plasmid were assembled using Gibson Assembly kit (New England Biolabs) to obtain the resulting genomic plasmid, pSAD-P-mCherry.

Recombinant virus was recovered as described previously [16]. Briefly, BSR-T7 cells (7 x 10^5^ cells) were transfected using Lipofectamine 2000 (Thermo Fischer Scientific) with 10 µg of genomic plasmid pSAD-P-mCherry, in addition to 5 µg pTit-N SAD, 2.5 µg pTit-P SAD and pTit-L SAD, 1.25 µg pTit-G SAD, which encode the N, P, L and G proteins of RABV strain SAD L16-B19 y. Nine days post-transfection, the supernatant was passed on fresh BSR cells, and the recombinant viruses were amplified for three days and detected by the fluorescence of the P-mCherry protein. The pre-stock of amplified viruses was titrated by IF staining with anti-N antibody. This pre-stock was used to infected BSR cells (MOI = 0.1). Four days post-infection, the supernatant was collected, aliquoted and stored at -80°C for further experiments. The P-mCherry fusion gene was sequenced and revealed no mutations.

## Antibodies and drugs

The rabbit polyclonal anti-P antibody was previously described [8] and was used at a 1:2000 dilution. Mouse monoclonal antibodies against RPS16 (Abcam, ab155961), RPS2 (Abcam, ab177951), and RPL40 (Abcam, ab109227) were used at dilutions of 1:150, 1:150, and 1:375, respectively. Rabbit polyclonal antibodies against RPS6, RPL18, and RPL27A (Atlas Antibodies; HPA031153, HPA046572, and HPA063379) were used at dilutions of 1:150, 1:300, and 1:150, respectively. A rabbit polyclonal antibody against RPL29 (Proteintech, 15799-1-AP) was used at a dilution of 1:400. Mouse monoclonal antibodies against eIF4G and eIF4H (Cell Signaling Technology; 2461 and 3469) were used at dilutions of 1:200 and 1:400, respectively. A rabbit polyclonal antibody against RPL39 (Sigma-Aldrich, SAB4502671) was used at a dilution of 1:600. A mouse monoclonal antibody against G3BP1 (Sigma-Aldrich, WH0010146M1) was used at a dilution of 1:50. An Alexa Fluor™ 488- conjugated anti-puromycin antibody (Sigma-Aldrich, MABE343-AF488) was used at a dilution of 1:250. Alexa Fluor 488-, 568-, or 647-conjugated secondary antibodies were purchased from Molecular Probes and used at a 1:1000 dilution. Puromycin (PYM), Emetine, Anisomycin, Cycloheximide (CHX), Nocodazole (NCZ) and Cytochalasin D (CytD) were from Sigma.

## Viral infection

Monolayers of cells (80% of confluence) were infected with the indicated virus at a multiplicity of infection (MOI) of 3, diluted in DMEM. After 1h of incubation at room temperature (RT), the inoculum was removed and the cells were maintained in DMEM supplemented with 2.5% FCS and incubated at 37°C and 5%CO_2_. For “mock” conditions, cells were incubated 1h at RT in DMEM only.

## Ribopuromycylation

Monolayers of cells were seeded on glass coverslips in DMEM supplemented with 10% FCS, 100 U/mL penicillin, and 100 µg/mL streptomycin at a density of 2 × 10⁵ cells per well. The following day, cells were pretreated or not with anisomycin (9.4 µM) for 15 min before infection. At 16 hours post-infection (h.p.i.) at 37°C and 5%CO_2_, cells were treated with 90 µM puromycin and 45 µM emetine for 5 minutes at 37°C then immediately placed on ice and rinsed once with cold phosphate buffer saline (PBS). Cells were then fixed for 10 min at -20°C with pre-chilled 37.5% methanol and 50% acetone before rehydration for 3 min in PBS. Cells were maintained at 4°C in PBS prior immunofluorescence staining.

## Immunofluorescence staining and confocal microscopy

After saturation with 1% Bovine Serum Albumin (BSA) in PBS, cells were incubated with the indicated primary antibodies for 1h at RT, washed with PBS and incubated for 45 min with Alexa fluor conjugated secondary antibodies. Cells were then incubated for 5 min with 0.1 mg/mL DAPI in PBS (to stain the nuclei) before washing and mounting with Immu-Mount (Thermo Fisher Scientific). Images were sequentially acquired using a laser scanning SP8 confocal microscope (Leica) with a 63x oil immersion objective (HC Plan Apo, NA: 1.4, Leica). Image analysis was performed using ImageJ software.

## Homopropargylglycine (HPG) pulse-labeling and click chemistry

Cells on coverslips were mock-infected or infected with RABV (MOI = 3). At times post-infection indicated, medium was removed and replaced with L-methionine-free DMEM containing 2% FBS for 1h to deplete methionine prior to the addition of HPG (Molecular Probes) at a final concentration of 1 mM for the indicated time. As control, CHX (10µM) was included in presence of HPG to block de novo protein synthesis. Cells were then placed on ice and rinsed once with cold PBS. After fixation for 20 min with 4% PFA (paraformaldehyde) and permeabilization for 5 min with 0.1% Triton X-100 in PBS, cells were processed for IF staining (see above). Samples were then subjected to click reaction using the Click-iT™ Plus Alexa Fluor™ 488 Picolyl Azide Toolkit (ThermoFisher Scientific). After removal of the reaction cocktail, cells were washed with PBS and mounted on slides.

## Statistical analysis

Data are expressed as mean ± standard deviation (SD) and statistical analysis was performed using Prism (GraphPad 10 software). Data comparison was performed using the Student *t*-test. Statistical details of each experiment are indicated in the figure legends.

## Results

### Detection of ribopuromycylated products in NBs

RABV mRNA transcription takes place inside NBs [17]. These viral mRNAs are structurally similar to cellular mRNAs and are therefore efficiently translated by the ribosomes. Whether translation of viral mRNAs occurs in NB and/or in the cytoplasm is unknown and a coupling of transcription and translation activities cannot be ruled out. To investigate this, we used ribopuromycylation (RPM) to map active translation sites in RABV infected cells [18]. In this method, cells are briefly treated with emetine, an elongation inhibitor, to immobilize actively translating 80S ribosomes on mRNAs, followed by a brief exposure to puromycin (PMY), a Tyr-tRNA mimetic. PMY is incorporated into the ribosome-associated nascent polypeptide chains, and the subcellular location of ribosome-associated nascent chains can then be detected by immunofluorescence (IF) using a monoclonal antibody against PMY. Due to the liquid nature of NBs, we had to adapt the labelling protocol described in the literature [18,19], as treatment with digitonin, used to release free PMY after cell fixation with PFA, led to the disappearance of NBs **(Figure 1)**. Instead, the cells were fixed in methanol/acetone, which allows both fixation and permeabilization. Under these conditions, the NBs are preserved and the free PMY molecules are eliminated during washing prior to immunolabelling (**Figure 1**). As expected, puromycin (PMY) signals were intense and uniformly distributed in all mock infected cells (**Figure 2**), indicating active protein synthesis. In RABV infected cells (16 h.p.i.), PMY signals remained evenly distributed within the cytoplasm and some signal was also detected in NBs in both U373-MG and BSR cells (**Figure 2**). Importantly, as shown in BSR cells (**Figure 2b**), anisomycin, a competitive inhibitor of PMY binding to ribosomes, prevents puromycylation both in the cytosol and in NBs. This inhibition confirms that the signals observed in NBs likely arise from *bona fide* ribosome-associated puromycylation, rather than non-specific accumulation of PMY. Moreover, the comparable sensitivity of cytosolic and NB-localized translation to anisomycin suggests that the translational machinery operates in both compartments.

**Figure 1.**
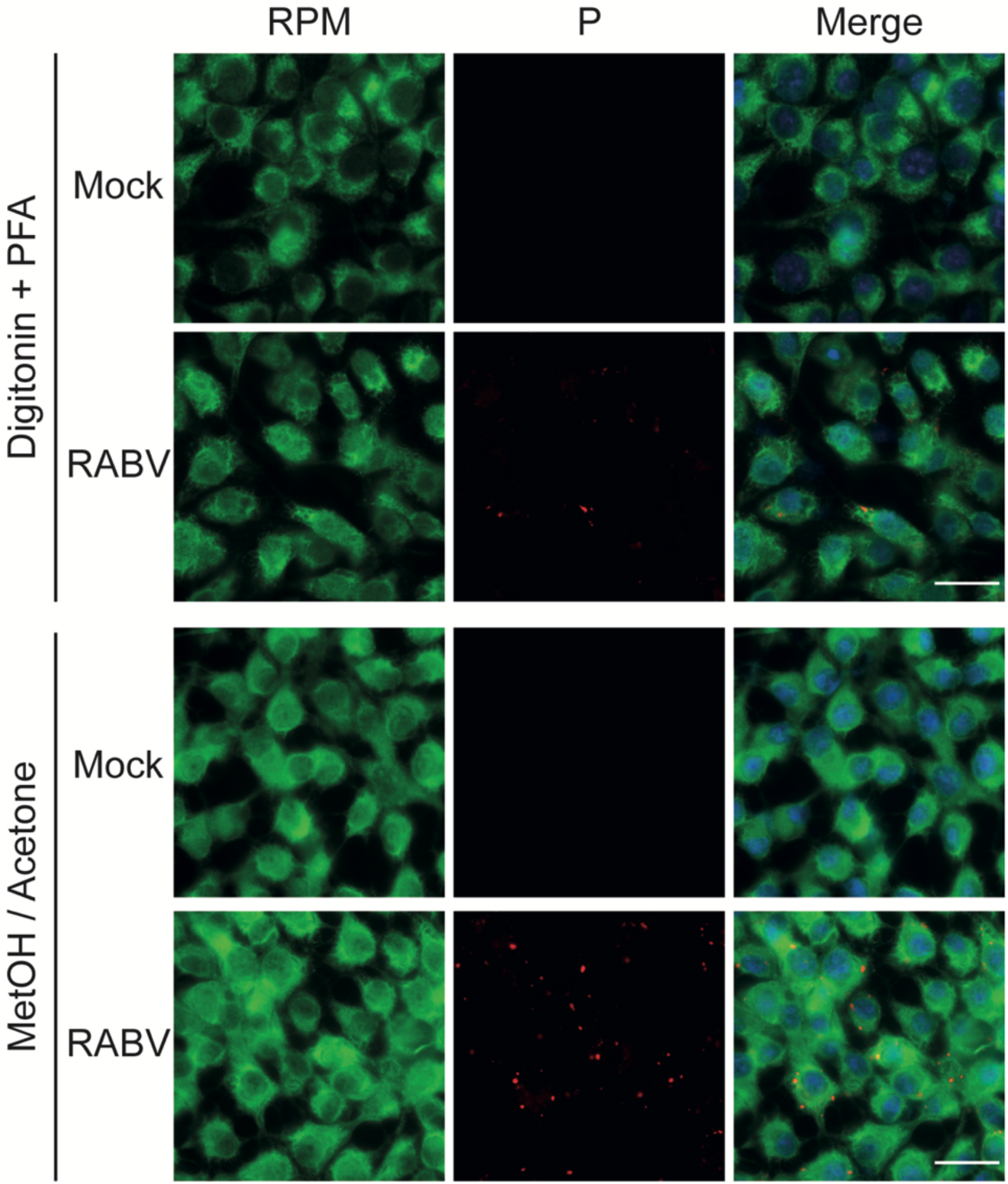
Detection of active ribosomes by RPM. BSR cells were infected with RABV virus at MOI 3. At 16 h.p.i, cells were incubated for 5 min at 37°C with PMY and emetine and, then either permeabilized with digitonin (0.015%) prior to fixation with 4% PFA (upper panel) or fixed with Methanol/Acetone (lower panel). After fixation, cells were subjected to anti-puromycin (green) and anti-P (red) immunostaining. Nuclei were stained with DAPI. Observations were made using an epifluorescence microscope (63x magnification). Scale bar: 40 μm.

**Figure 2.**
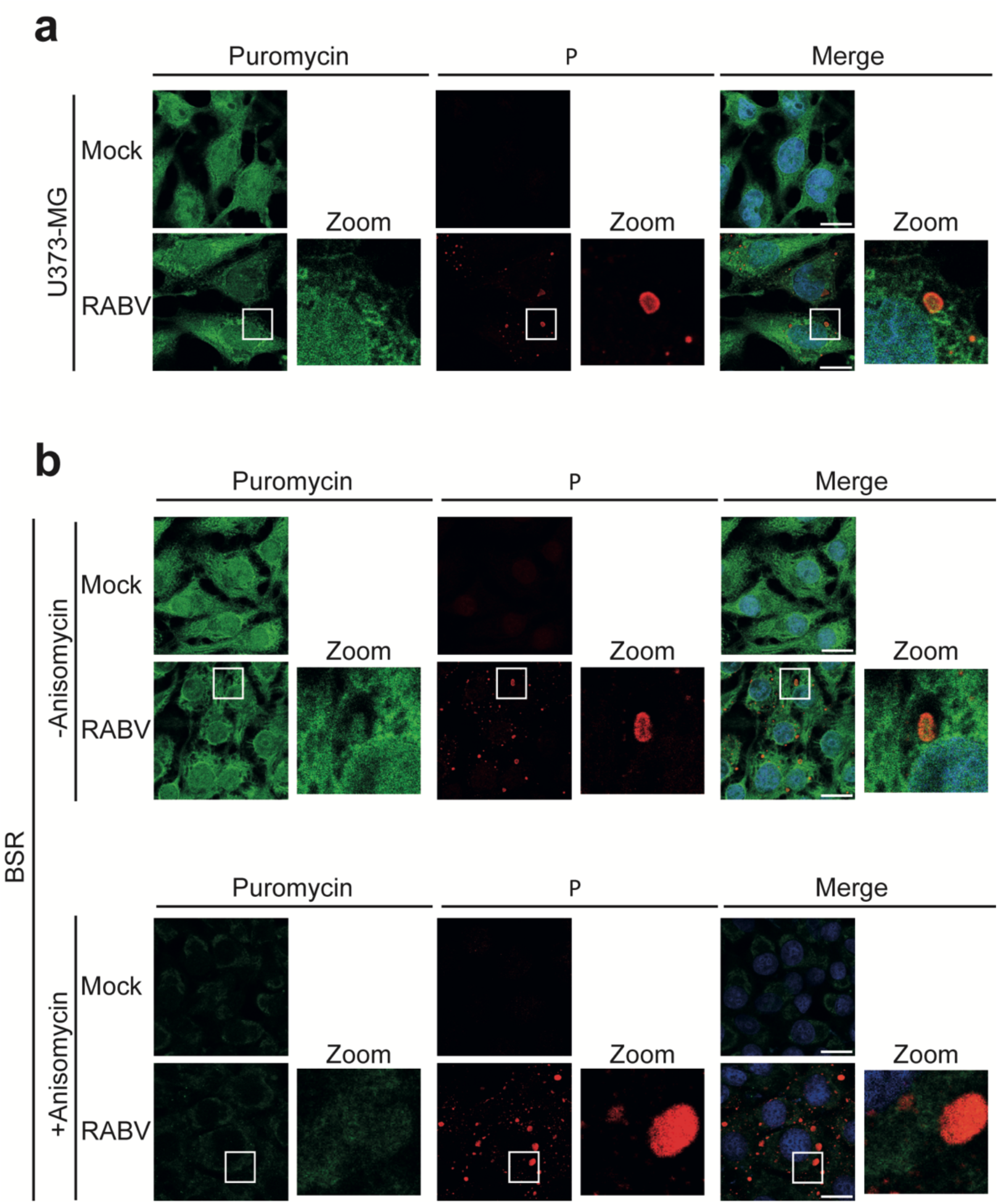
RPM labeling of RABV infected cells. U373-MG (upper panels) and BSR (lower panels) cells were infected or not infected (Mock) with RABV (MOI 3). At 16 h.p.i., cells were pretreated or not with anisomycin, as indicated, and then incubated for 5 min at 37°C with PMY and emetine. Cells were then fixed in 4% PFA and subjected to co-immunolabelling with anti-puromycin (green) and anti-P (red) antibodies. Nuclei were stained with DAPI. Images were acquired using a confocal microscope (63x magnification). Square boxes in the merge are magnified (zooms). Scale bar: 15μm.

## Ribosomal proteins and translation factors are excluded from NBs

The presence of viral mRNA molecules within NBs, together with the positive RPM labeling, suggest that viral translation may take place inside the viral factories. To address this question, we explored the localization of components of the cellular translation machinery in infected cells using IF microscopy (**Figure 3**). Protein components of both 40S (RPS6 and RPS16) (**Figure 3a**) and 60S (RPL18 and RPL27A) (**Figure 3b**) ribosomal subunits were found throughout cytoplasm but excluded from NBs. The same observation was made for the RPL40 protein (**Figure 3b**), constituent of the large ribosomal subunit, and required for replication of several MNV, as well as for cellular proteins involved in translation initiation, the Poly(A)binding protein (PABP) and eukaryotic translation initiation factors 4G (eIF4G) and 4H (eIF4H) (**Figure 3c**). These exclusions strongly suggest that active translation is unlikely to occur inside NBs, indicating that viral mRNAs are exported from viral factories to be translated in the cytoplasm.

**Figure 3.**
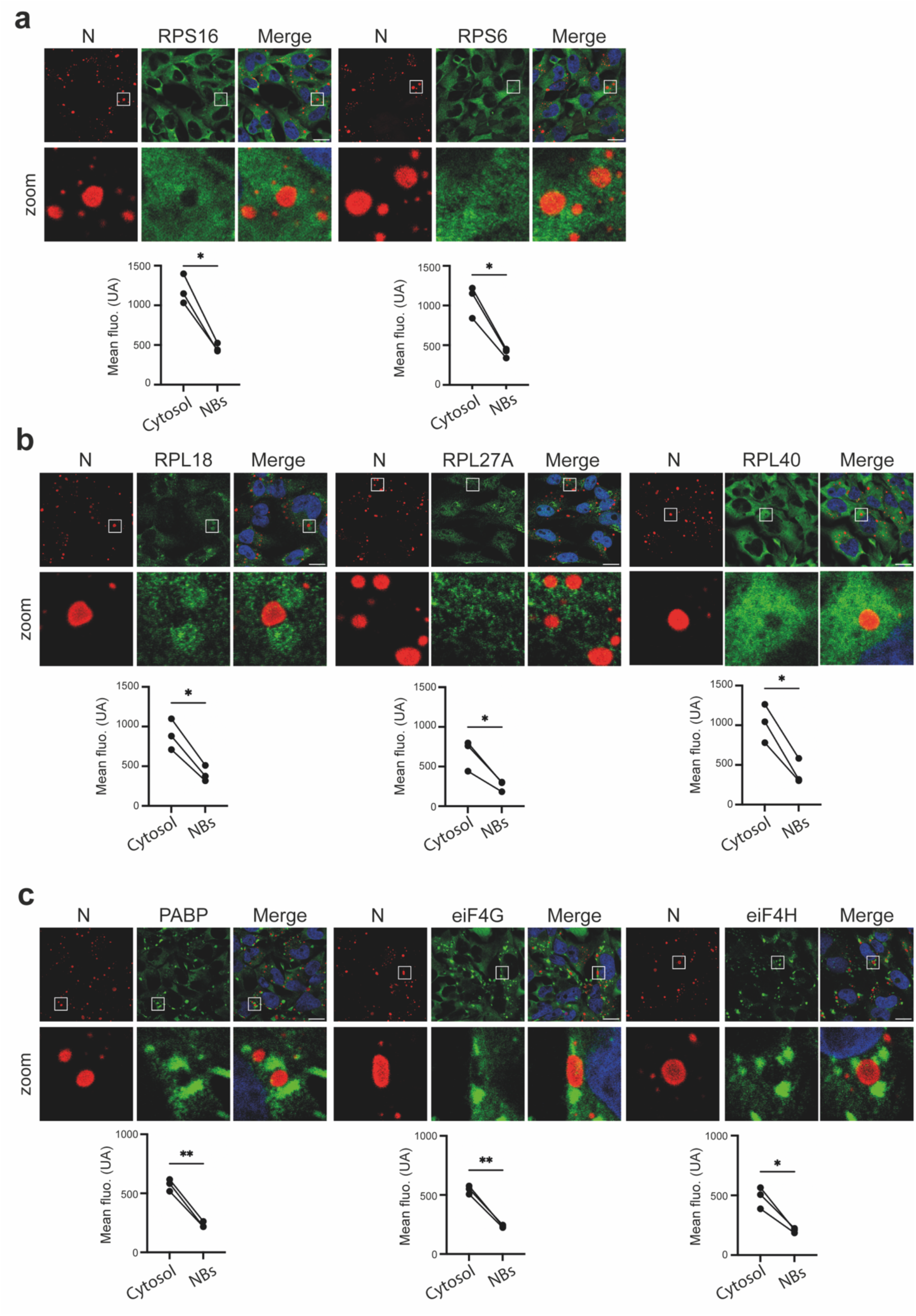
Ribosomal proteins and translation factors are excluded from NBs. U373-MG cells were infected with RABV (MOI 3). At 16 h.p.i., cells were fixed in 4% PFA and co- immunostained with anti-N antibody (red) and antibodies against ribosomal subunit proteins or translation factors, as indicated (green). Nuclei were stained with DAPI. Images were acquired using a confocal microscope (63x magnification). Square boxes are magnified (zooms). Scale bars: 15μm. Green fluorescence was measured in the cytosol and within NBs. For each category (cytosol and NB), 30–60 locations were quantified per experiment, corresponding to 10–20 cells. Each point represents the mean of 30–60 measurements from one independent experiment (n = 3), with paired measurements connected by lines. Statistical significance was assessed using a two-tailed paired t- test. (ns =not significant; *P < 0.05; **P < 0.01).

## Rapid concentration of newly synthesized proteins in NBs

To gain more insight into the dynamics of cellular and viral mRNA translation, newly synthesized proteins can be labeled with the methionine analogue homopropargylglycine (HPG) and subsequently conjugated to an azide fluorophore (Alexa Fluor 488-azide in the present case). This bioorthogonal labeling strategy permits quantitative and spatial assessment of the cellular “translatome” by fluorescence microscopy. We first pulse-labeled cells with HPG for 15 min and performed fluorescent microscopy analysis after cell counterstaining with anti-P antibody to confirm RABV infection and visualize the NBs (**Figure 4a**). In mock-infected cells, an abundant accumulation in the nucleus and a diffuse pattern in the cytoplasm were observed, in line with previous observations made by others [20][21]. In infected cells (16 h.p.i.), weak labelling in the cytosol and massive incorporation of HPG into the nucleus were also observed, as well as an accumulation of newly synthesized proteins in spherical clusters in the cytosol that corresponded to NBs as revealed by anti-P labeling. The nearly complete inhibition observed with cycloheximide (CHX) treatment confirms that HPG incorporation reflects *de novo* protein synthesis with no contribution from free HPG and that the spatial distribution observed corresponds to that of newly synthesized proteins. To refine the spatiotemporal analysis of protein synthesis in RABV-infected cells, HPG pulse-labeling was reduced to 5, 3, and 1 minute. Fluorescence levels were similar in the cytosol and NBs after a one-minute pulse **(Figure 4b)**. However, the mean NB-to-cytosol fluorescence ratio increased progressively with HPG pulse duration reaching 1.6 at 3 min and 2 at 5 min **(Figure 4b)**. Although this approach does not allow discrimination between host-derived and viral proteins, the results reveal a rapid concentration of newly synthesized proteins in NBs.

**Figure 4.**
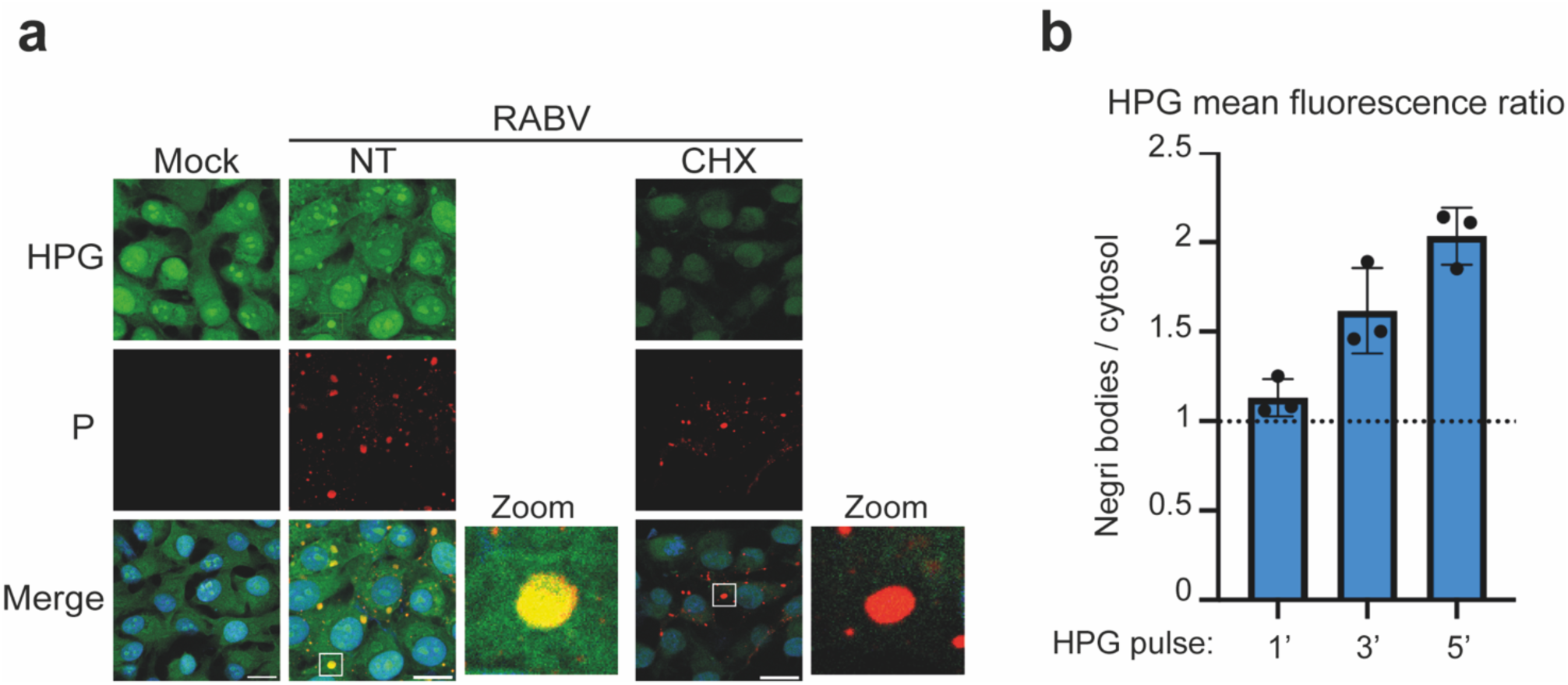
Newly synthesized proteins accumulate in NB. (a) BSR cells were infected or not (Mock) with RABV (MOI 3) and pulse-labeled with HPG for 15 min at 16 h.p.i. As control, BSR cells were pulse-labeled with HPG for 15 min in presence of CHX (10µM). Cells were then subjected to co-labeling to detect newly synthesized proteins (green) and the RABV phosphoprotein P (red). Nuclei were stained with DAPI. Images were acquired using a confocal microscope (63x magnification). Square boxes in the merge are magnified (zooms). Scale bars: 15μm. (b) BSR cells were infected with RABV (MOI 3) and pulse-labeled with HPG for 1, 3, or 5 min at 16 h.p.i. Cells were then analyzed as in (a), and HPG mean fluorescence intensity was measured in NBs and the cytosol using ImageJ software. For each condition, 40–60 measurements were performed for each category (NBs and cytosol). The graph shows the ratio of HPG fluorescence intensity between NBs and the cytosol (Negri bodies/cytosol). Each dot represents the ratio obtained from one independent experiment (n=3). Bars indicate the standard deviation (SD), and the dashed line indicates a ratio of 1, corresponding to equal HPG fluorescence intensity in NBs and the cytosol.

## Rapid concentration of newly synthesized proteins in viral factories is independent of tubulin and actin cytoskeleton

Given the absence of translation machinery in NBs and the rapid accumulation of newly synthesized proteins observed in these compartments, it was reasonable to think that the viral proteins resident in NBs are translated in the cytosol and then rapidly addressed to NBs.

To investigate the role of the cytoskeleton in the incorporation of newly synthesized proteins in NBs, we used various drugs which interfere with the architecture of the cytoskeleton.

To test the hypothesis that viral proteins are first synthesized outside NBs and then being transported into them by a microtubule (MT)-dependent transport, BSR cells were infected with RABV and treated for 2 hours with 10 μM nocodazole (NCZ) before HPG pulse-labelling for 3, 5 or 15 min. In the presence of NCZ, the microtubule network (assessed by IF analysis with anti-tubulin labelling) is effectively depolymerized, both in infected and mock-infected conditions (**Figure 5a, lower panel**). By contrast, NCZ treatment does not affect protein translation or protein concentration in NBs as incorporation of HPG was similar to that observed in mock-treated cells (**Figure 5a**). Indeed, quantification of fluorescence intensity reveals no significant difference between the NBs of cells treated or untreated with NCZ (**Figure 5b**) and confirmed the early, rapid and massive concentration of newly translated protein into NBs.

**Figure 5.**
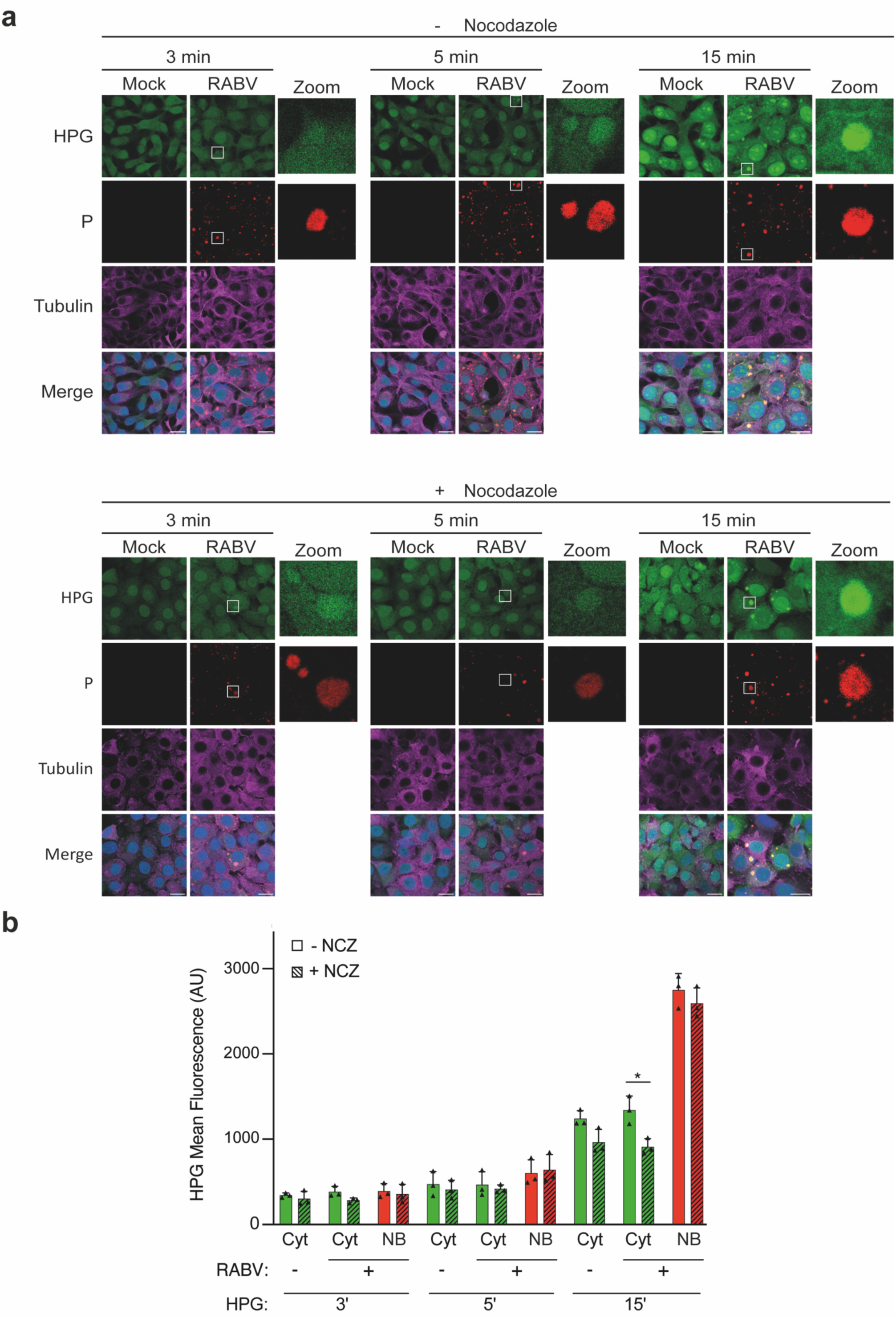
Impact of microtubule network disruption on the localization of newly synthesized proteins in NBs. (a) BSR cells were infected or not (Mock) with RABV (MOI 3) and pulse-labeled with HPG for 3, 5 min or 15 min at 16 h.p.i. Prior to HPG pulse cells were pretreated (lowel panels) or not (upper panels) with nocodazole (NCZ) for 2 hours at 37°C. Cells were then subjected to co-labeling to detect newly synthesized proteins (green), the RABV phosphoprotein P (red) and tubulin (magenta). Nuclei were stained with DAPI. Images were acquired using a confocal microscope (63x magnification). Scale bar: 15μm. (b) HPG mean fluorescence intensity was measured in NBs and the cytosol using ImageJ software. For each category, 30–60 locations were quantified, corresponding to 10–20 cells per experiment. The bars represent the mean values and standard deviations of three independent experiments; ns = not significant; *P < 0.05 (Student’s *t*-test).

Similarly, in order to determine whether the actin cytoskeleton was involved in the rapid concentration of *de novo* synthesized proteins into NBs, BSR cells were infected with RABV and treated for 2 hours with Cytochalasin D (Cyto D) before HPG pulse-labelling for 3, 5 or 15 min. Cyto D treatment, which dirupts actin polymerization (as verified by phalloidin labelling, **Figure 6a**), results in an increased fragmentation of NBs as previously described (**Figure 6b**). As in absence of MT network, the disruption of actin filaments does not affect protein translation and concentration in NBs (**Figure 6b**) and the kinetics of concentration of newly synthesized proteins into NBs is unchanged in treated or mock-treated cells (**Figure 6c**).

**Figure 6.**
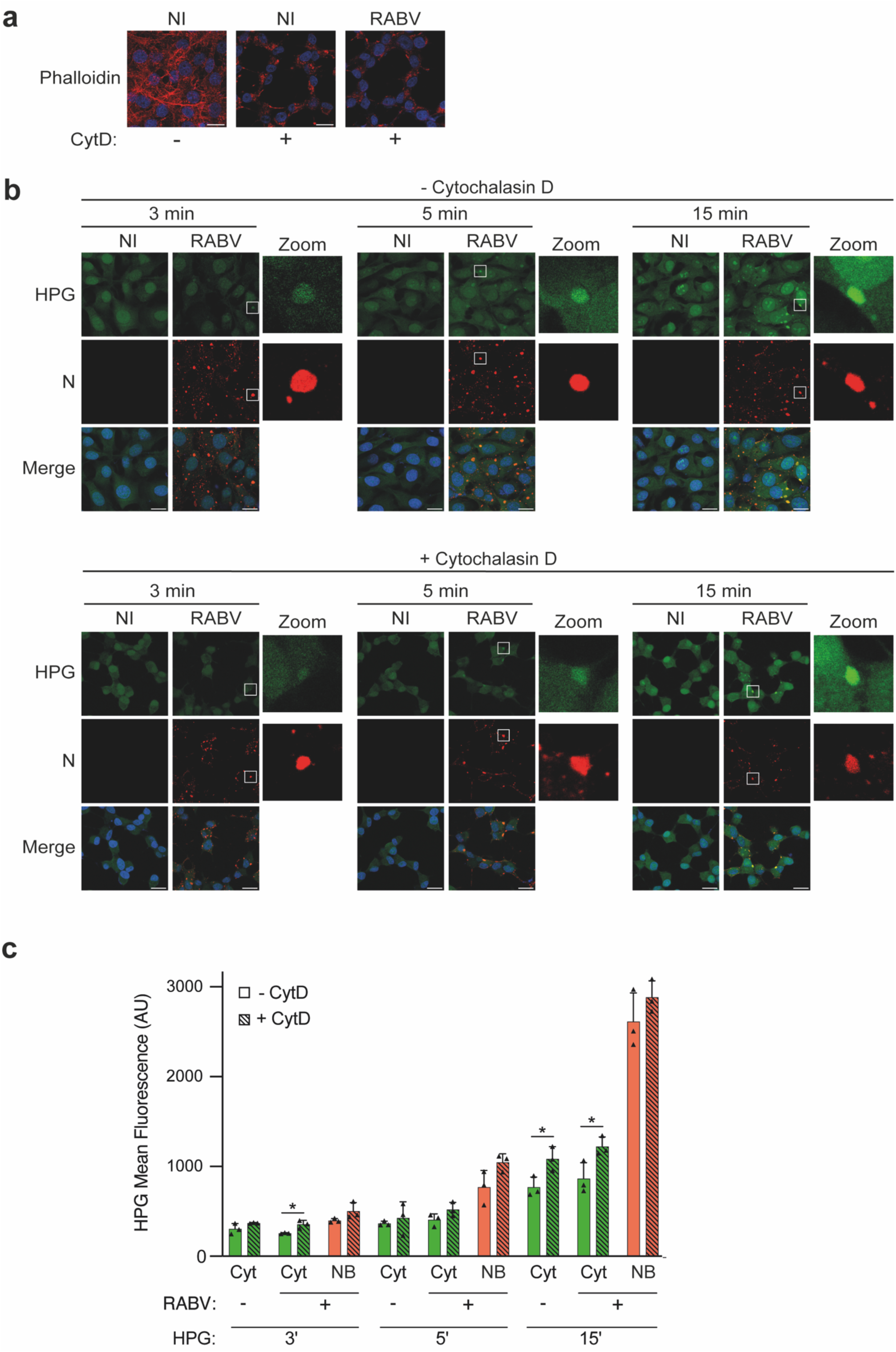
Impact of actin network disruption on the localization of newly synthesized proteins in NBs. (a) BSR cells were infected with RABV (MOI 3) or left non-infected (Mock) and treated or not with cytochalasin D (CytD) for 2 h at 37 °C at 14 h.p.i. Then, cells were fixed with 4% PFA and stained with phalloidin (red). Nuclei were stained with DAPI. Images were acquired using a confocal microscope (63x magnification). Scale bar: 15μm. (b) BSR cells were infected or not (Mock) with RABV (MOI 3) and pulse-labeled with HPG for 3, 5 min or 15 min at 16 h.p.i. Prior to HPG pulse cells were pretreated (lower panels) or not (upper panels) with cytochalasin D (NCZ) for 2 hours at 37°C. Cells were then subjected to co-labeling to detect newly synthesized proteins (green), the RABV phosphoprotein N (red). Nuclei were stained with DAPI. Images were acquired using a confocal microscope (63x magnification). Scale bar: 15μm. (c) HPG mean fluorescence intensity was measured in NBs and the cytosol using ImageJ software. For each category, 30–60 locations were quantified, corresponding to 10–20 cells per experiment. The bars represent the mean values and standard deviations of three independent experiments; *P < 0.05, (Student’s *t*-test).

Taken together, these results demonstrate that neither the MT network nor actin filaments are essential for the rapid concentration of newly translated proteins in viral factories. This suggests that most proteins residing in NBs, at least the viral N, P, and L proteins, could be translated at the periphery of NBs to be rapidly incorporated after translation without cytoskeleton-dependent transport.

## RABV infection triggers SGs formation

As previously described [22], in RABV infected cells we could observed the formation of SGs, as revealed by the detection of SG-marker proteins Ras GTPase-activating protein-binding protein 1 (G3BP1), some of which in contact with NBs. This observation was made in U373-MG cells (**Figure 7a**), but not in BSR cells, which are highly permissive to viral infection due to deficiencies in the RIG-I pathway [23] and low levels of activatable PKR [24]. Of note, in U373-MG infected cells displaying SGs, the RPM signal was markedly reduced (**Figure 7a**). Similarly, in these cells the HPG signal was undetectable in both the cytosol and NBs, whereas it was clearly present, and concentrated in NBs, in infected but non-stressed cells (**Figure 7b**). These observations are consistent with published data showing that SG formation is associated with translational repression and polysome disassembly [25][26].

**Figure 7.**
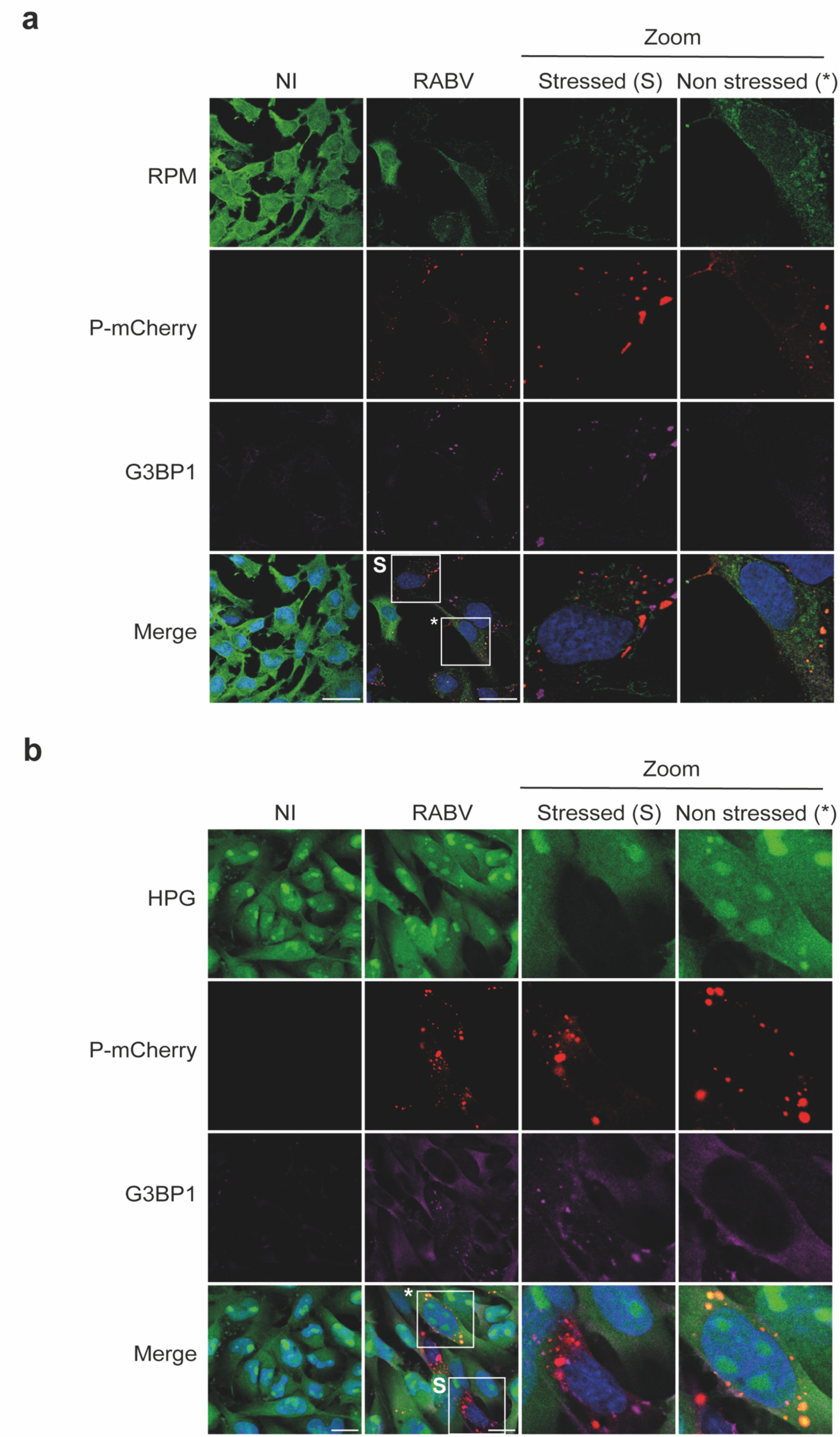
Translation is arrested in RABV infected cells containing SGs. (a) U373-MG cells were infected with RABV-PmCherry (MOI 3). At 16 h.p.i., cells were incubated for 5 min at 37°C with PMY and emetine. Cells were then fixed in 4% PFA and subjected to co-immunolabelling with anti-puromycin (green) and anti-G3BP (magenta) antibodies. (b) U373-MG cells were infected with RABV-PmCherry (MOI 3) and pulse-labeled with HPG for 5 min at 16 h.p.i. Cells were then subjected to co-labeling to detect newly synthesized proteins (green) and G3BP (magenta). In (a) and (b), nuclei were stained with DAPI. Images were acquired using a confocal microscope (63x magnification). Square boxes in the merge are magnified; S, stressed cell; asterisks, non-stressed cell. Scale bar: 15μm.

## Some ribosomal proteins and translation factors are concentrated in SGs

To further investigate the translation events in stressed cells upon RABV infection, we examined the distribution of cellular translation components in these cells (**Figure 8**). In stressed cells, protein components of 40S ribosomal subunit (RPS6 and RPS16) were found throughout cytoplasm and significantly concentrated in SGs (**Figure 8a**). The same observation was made for cellular proteins involved in translation initiation, the Poly(A)binding protein (PABP) and eukaryotic translation initiation factors 4G (eIF4G) and 4H (eIF4H) (**Figure 8b**). On the contrary, except for RPL18, protein components of the 60S ribosomal subunit (RPL27A and RPL40) were found equally in the cytosol and SG (**Figure 8c**). Overall, these observations are consistent with the known characteristics of SGs that contain mRNAs associated with translation initiation complexes (i.e., small 40S ribosomal subunits), but not the large ribosomal subunit components.

**Figure 8.**
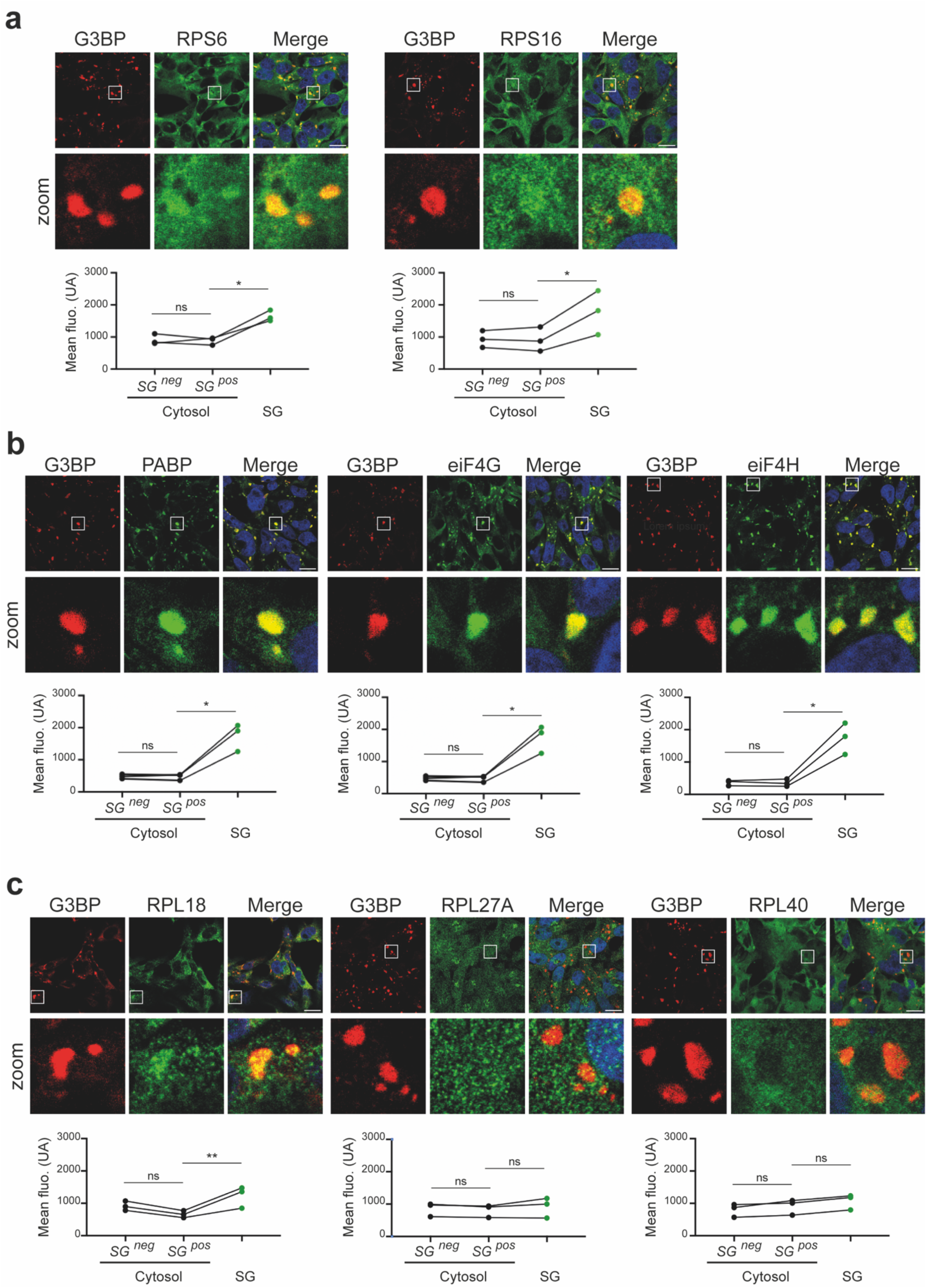
RABV infection promotes the enrichment of 40S ribosomal proteins and translation initiation factors in SGs. U373-MG cells were infected with RABV (MOI 3). At 16 h.p.i., cells were fixed in 4% PFA and co- immunostained with anti-G3BP antibody (red) and antibodies against ribosomal subunit proteins or translation factors, as indicated (green). Nuclei were stained with DAPI. Images were acquired using a confocal microscope (63x magnification). Scale bars: 15μm. Graphs show mean fluorescence intensity (green channel) in the cytosol of non-stressed cells (without SG), the cytosol of stressed cells (with SG), and within SG. For each category, 30–60 locations were quantified, corresponding to 10–20 cells per experiment. Each point represents one independent experiment (n = 3), with paired measurements connected by lines. Statistical analysis was performed using a two-tailed paired *t*-test (ns = not significant; *P < 0.05; **P < 0.01).

## Discussion

### Spatial separation of transcription and translation in RABV-infected cells

In this study, we investigated the existence of specific sites for the translation of viral transcripts in RABV infected cells. In particular, the high concentration of viral mRNAs within NBs could suggest that their translation could take place in this compartment or in its immediate vicinity. The absence of essential ribosomal components inside NBs strongly argues against this and indicates that translation events do not occur within the viral factories. The fact that a weak positive signal is detected in the NBs with RPM is probably due to a technical artifact because, to preserve the integrity of the NBs, we had to adjust the protocol by avoiding permeabilization with digitonin. Under these conditions, the methanol/acetone fixation method may not be efficient to remove free RPM, which could be trapped in the NBs and cause a residual positive signal. The most parsimonious interpretation of our data supports a model in which viral mRNAs are exported from the NBs into the cytoplasm for translation and then, newly synthesized proteins accumulate in the viral factories. This « export model » brings new insights for understanding the functional architecture of NBs. Rather than functioning as fully autonomous compartments encompassing all steps of viral gene expression, NBs appear to present selective permeability that allows mRNA export while excluding the translation machinery. Such selective permeability likely relies on their liquid nature based on LLPS. The molecular mechanisms governing mRNA export from NBs remain to be elucidated but may involve specific viral or host factors that facilitate mRNA release into the cytosol. In this regard, a study of Ebola virus identified the host mRNA export factor NXF1, recruited by the viral nucleoprotein NP, as a potential mediator of mRNA trafficking from viral inclusion bodies [27], raising the possibility that similar mechanisms may operate in rabies virus NBs.

## Rapid protein concentration in viral factories

The bioorthogonal labeling experiments using HPG revealed the rapid concentration of newly synthesized proteins in NBs. This observation indicates that following their cytoplasmic translation, viral proteins are efficiently and quickly recruited in the viral factories. This could also be the case for some cellular proteins recruited in NBs.

Importantly, treatments with nocodazole, which depolymerizes microtubules, or with cytochalasin D, which depolymerizes actin filaments, do not impact this recruitment process, demonstrating that neither microtubules nor actin filaments are essential for this rapid protein concentration. These observations suggest a mechanism of « diffusion-capture », where proteins freely diffuse through the cytoplasm and are selectively retained within NBs through specific molecular interactions with components of NBs. This « diffusion-capture » model is particularly attractive given the liquid properties of NBs and their capacity for selective molecular partitioning based on LLPS. However, this does not rule out local translation in the immediate vicinity of NBs, which could further minimize the distance proteins must travel to reach the viral factory. The two mechanisms could therefore potentially operate in concert.

## Translation dynamics in stress granule-positive infected cells

Our results show that some ribosomal proteins and translation factors relocalize to stress granules (SGs) in stressed cells infected with RABV. SGs are generally associated with translational repression and antiviral responses. Indeed, in RABV-infected U373-MG cells that contains SGs, translation is arrested as observed in our bioorthogonal labeling experiments. However, RABV replication is maintained in U373MG. The ability of RABV to sustain infection in cells competent for SG formation suggests that the virus has evolved mechanisms to maintain viral protein synthesis via SG-independent pathways. Along this line, studies have shown that the localization of mRNA in SGs is compatible with translation [28]. Thus, viral mRNAs exported from NBs to SGs could access the translation machinery associated with SGs. Alternatively, a subset of translation-competent ribosomes could remain outside of SGs, or viral mRNAs may possess features that allow efficient translation when host protein synthesis is reduced as suggested by Lee et al [15]. Finally, SGs formed during a viral infection are extremely dynamic, appearing and then disappearing suddenly in the cytoplasm [22]. Therefore, we cannot exclude out the possibility that viral protein translation occurs during SG-free phases. Further studies investigating the interactions between viral mRNAs, SG components, and ribosomes will be important to clarify the mechanisms that support viral protein synthesis in stressed cells.

## Relationships between translation and viral factories across *Mononegavirales*

The organizational principles we have uncovered for NBs show both similarities and differences compared to viral factories of other RNA viruses. Many negative-sense RNA viruses, including measles virus [12,29], respiratory syncytial virus [13,30], and Ebola virus [14,31], form cytoplasmic inclusion bodies that serve as replication and transcription centers. Even though it has been shown that orthopneumoviruses sequester various components of the translational preinitiation complex machinery into viral inclusion bodies [32], similar to our findings with NBs, these viral factories typically exclude ribosomes, suggesting that mRNA export for cytoplasmic translation may be a common feature of negative-sense RNA virus replication strategies. The spatial separation of transcription and translation observed for RABV may provide advantages such as protecting viral mRNAs from degradation or allowing for post-transcriptional processing. Moreover, the exclusion of translation machinery from NBs may help to maintain a specialized environment required for genome replication and transcription, preventing interference between these processes and translation.

## Limitations and technical considerations

Several limitations of our study should be acknowledged. First, while the HPG labeling approach effectively highlights newly synthesized proteins, it cannot distinguish between viral and host proteins. The rapid concentration we observe in NBs likely reflects primarily viral proteins, given the known composition of these structures, but contributions from host proteins cannot be excluded. Future studies using more selective labeling strategies or proteomics approaches could provide greater resolution of the protein composition dynamics in NBs. Second, the experiments conducted with Nocodazole and Cytochalasin D ruled out major roles for microtubules and actin filaments but did not extensively test all potential mechanisms of protein trafficking. Other cytoskeletal elements or chaperones could contribute to protein concentration in NBs. Furthermore, the role of liquid-liquid separation of NBs, which we propose as a potential mechanism for protein concentration, requires direct biophysical characterization to be definitively established.

## Conclusions

The exclusion of ribosomal proteins and translation factors from NBs, combined with the rapid, cytoskeleton-independent concentration of newly synthesized proteins in these structures, supports a model of spatial separation between transcription and translation in the case of RABV infection. This organization likely optimizes viral replication by exploiting cytoplasmic translation resources while maintaining a specialized environment dedicated to genome replication and transcription in NBs. Our findings contribute to a better understanding of how viruses manipulate cellular spatial organization to facilitate their replication cycles and suggest new perspectives on the functional architecture of viral factories.

## Data availability statement

The datasets used and/or analyzed during the current study are available from the corresponding author on reasonable request.

## Acknowledgments

This work was supported by the « Prix Bettencourt Coups d’élan pour la recherche française » attributed to YG, a grant from the Agence Nationale de la Recherche, France, (ANR CE11, LiquidFact) and a grant from the Fondation pour la Recherche Médicale, France, to Y.G. The present work has benefited from Imagerie-Gif core facility supported by I’Agence Nationale de la Recherche (ANR-11-EQPX-0029/Morphoscope, ANR-10-INBS-04/FranceBioImaging; ANR-11- IDEX-0003-02/ Saclay Plant Sciences).

## Notes

### Competing Interest Statement

The authors have declared no competing interest.

## References

1. Rozman, B., Fisher, T. & Stern-Ginossar, N. Translation—A tug of war during viral infection. Molecular Cell 83, 481–495 (2023).

2. Pakos-Zebrucka, K., et al. The integrated stress response. EMBO Rep 17, 1374–1395 (2016).

3. Hofmann, S., Kedersha, N., Anderson, P. & Ivanov, P. Molecular mechanisms of stress granule assembly and disassembly. Biochimica et Biophysica Acta (BBA) - Molecular Cell Research 1868, 118876 (2021).

4. Riggs, C. L., Kedersha, N., Ivanov, P. & Anderson, P. Mammalian stress granules and P bodies at a glance. J Cell Sci 133, jcs242487 (2020).

5. Brownsword, M. J. & Locker, N. A little less aggregation a little more replication: Viral manipulation of stress granules. WIREs RNA 14, e1741 (2023).

6. Hampson, K. et al. Estimating the Global Burden of Endemic Canine Rabies. PLOS Neglected Tropical Diseases 9, e0003709 (2015).

7. Albertini, A. A. V., Ruigrok, R. W. H. & Blondel, D. Chapter 1 - Rabies Virus Transcription and Replication. in Advances in Virus Research (ed. Jackson, A. C.) vol. 79 1–22 (Academic Press, 2011).

8. Lahaye, X. et al. Functional Characterization of Negri Bodies (NBs) in Rabies Virus-Infected Cells: Evidence that NBs Are Sites of Viral Transcription and Replication. J Virol 83, 7948–7958 (2009).

9. Nikolic, J. et al. Negri bodies are viral factories with properties of liquid organelles. Nat Commun 8, 58 (2017).

10. Heinrich, B. S., Maliga, Z., Stein, D. A., Hyman, A. A. & Whelan, S. P. J. Phase Transitions Drive the Formation of Vesicular Stomatitis Virus Replication Compartments. mBio 9, (2018).

11. Guseva, S. et al. Measles virus nucleo- and phosphoproteins form liquid-like phase-separated compartments that promote nucleocapsid assembly. Sci Adv 6, (2020).

12. Zhou, Y., Su, J. M., Samuel, C. E. & Ma, D. Measles Virus Forms Inclusion Bodies with Properties of Liquid Organelles. Journal of Virology 93, e00948–19 (2019).

13. Rincheval, V. et al. Functional organization of cytoplasmic inclusion bodies in cells infected by respiratory syncytial virus. Nature Communications 8, 563 (2017).

14. Hoenen, T. et al. Inclusion Bodies Are a Site of Ebolavirus Replication. J Virol 86, 11779–11788 (2012).

15. Lee, A. S.-Y., Burdeinick-Kerr, R. & Whelan, S. P. J. A ribosome-specialized translation initiation pathway is required for cap-dependent translation of vesicular stomatitis virus mRNAs. Proc. Natl. Acad. Sci. U.S.A. 110, 324–329 (2013).

16. Wirblich, C. & Schnell, M. J. Rabies Virus (RV) Glycoprotein Expression Levels Are Not Critical for Pathogenicity of RV. J Virol 85, 697–704 (2011).

17. Lahaye, X. et al. Functional Characterization of Negri Bodies (NBs) in Rabies Virus-Infected Cells: Evidence that NBs Are Sites of Viral Transcription and Replication. Journal of Virology 83, 7948–7958 (2009).

18. David, A. et al. Nuclear translation visualized by ribosome-bound nascent chain puromycylation. J Cell Biol 197, 45–57 (2012).

19. Bastide, A., Yewdell, J. W. & David, A. The RiboPuromycylation Method (RPM): an Immunofluorescence Technique to Map Translation Sites at the Sub-cellular Level. Bio Protoc 8, (2018).

20. Beatty, K. E. et al. Fluorescence Visualization of Newly Synthesized Proteins in Mammalian Cells. Angewandte Chemie International Edition 45, 7364–7367 (2006).

21. Teo, C. S. H., Serwa, R. A. & O’Hare, P. Spatial and Temporal Resolution of Global Protein Synthesis during HSV Infection Using Bioorthogonal Precursors and Click Chemistry. PLOS Pathogens 12, e1005927 (2016).

22. Nikolic, J., Civas, A., Lama, Z., Lagaudrière-Gesbert, C. & Blondel, D. Rabies Virus Infection Induces the Formation of Stress Granules Closely Connected to the Viral Factories. PLoS Pathog 12, e1005942 (2016).

23. Habjan, M., Penski, N., Spiegel, M. & Weber, F. T7 RNA polymerase-dependent and -independent systems for cDNA-based rescue of Rift Valley fever virus. Journal of General Virology 89, 2157–2166 (2008).

24. Langland, J. O. & Jacobs, B. L. The Role of the PKR-Inhibitory Genes, E3L and K3L, in Determining Vaccinia Virus Host Range. Virology 299, 133–141 (2002).

25. Kedersha, N. et al. Stress granules and processing bodies are dynamically linked sites of mRNP remodeling. J Cell Biol 169, 871–884 (2005).

26. Souquere, S. et al. Unravelling the ultrastructure of stress granules and associated P-bodies in human cells. Journal of Cell Science 122, 3619–3626 (2009).

27. Wendt, L. et al. Evidence for Viral mRNA Export from Ebola Virus Inclusion Bodies by the Nuclear RNA Export Factor NXF1. J Virol 96, e00900–22.

28. Mateju, D. et al. Single-Molecule Imaging Reveals Translation of mRNAs Localized to Stress Granules. Cell 183, 1801–1812.e13 (2020).

29. Guseva, S. et al. Structure, dynamics and phase separation of measles virus RNA replication machinery. Current Opinion in Virology 41, 59–67 (2020).

30. Galloux, M. et al. Minimal Elements Required for the Formation of Respiratory Syncytial Virus Cytoplasmic Inclusion Bodies *In Vivo* and *In Vitro*. mBio 11, e01202–20, /mbio/11/5/mBio.01202-20.atom (2020).

31. Bodmer, B. S. et al. Ebola virus inclusion bodies are liquid organelles whose formation is facilitated by nucleoprotein oligomerization. Emerging Microbes & Infections 12, 2223727 (2023).

32. Jobe, F. et al. Viral PIC-pocketing: RSV sequestration of translational preinitiation complexes into bi- phasic biomolecular condensates. J Virol 98, e00153–24 (2024).

